# Machine Learning Driven Simulations of the SARS-CoV-2 Fitness Landscape from Deep Mutational Scanning Experiments

**DOI:** 10.1101/2024.09.20.614179

**Authors:** Aleksander E. P. Durumeric, Sean McCarty, Jay Smith, Jonas Köhler, Katarina Elez, Lluís Raich, Patricia A. Suriana, Terra Sztain

## Abstract

Predicting protein variant effects is a key challenge in preparing for pathogenic viral strains, understanding mutation-linked diseases, and designing new proteins. Protein sequence-structure-function relationships are difficult to model due to complex allosteric and epistatic effects. To investigate efficient modeling strategies, we trained supervised machine learning (ML) models with deep mutational scanning (DMS) libraries of SARS-CoV-2 receptor binding domain (RBD) sequences labeled with angiotensin converting enzyme 2 (ACE2) binding affinity. These models demonstrate superior performance predicting combinatorial mutation effects compared to adding or averaging the effects of point mutations and exhibit strong extrapolative performance ranking omicron variants when training only on wild type (WT) variants. We characterize the RBD fitness landscape combining ML with Markov Chain Monte Carlo simulations to predict evolutionary patterns from the WT sequence, and generate comparable sequence profiles to high fitness sequences in DMS data predicting mutations in unseen omicron variants. These models provide insight into the relationship between RBD sequence elements, and offer a new perspective on the use of DMS to predict emerging viral strains, which we anticipate will be applicable to other evolutionary prediction tasks. To facilitate application and future development of this strategy, we introduce Mavenets: https://github.com/SztainLab/mavenets.

## Introduction

The SARS-CoV-2 virus, which led to the COVID-19 pandemic, continues to evolve, accumulating mutations that evade immune responses and therapies.^1–4^ Preparation for future variants remains challenging due to the large number of possible mutations that could cause virulent strains. Most problematic mutations occur in the receptor binding domain (RBD) of the spike protein, which binds angiotensin converting enzyme 2 (ACE2) on host cells as the first point of contact for viral infection.^5^ Starr et al. performed deep mutational scanning (DMS) on the spike RBD in 2020,^6^ providing sequence-function information characterizing the effect of 10^5^ RBD sequence variants on ACE2 binding, a strong indicator of viral fitness.^7–9^ Additional experiments initialized from the B1.351 variant in 2021^10^ and omicron BA.1 and BA.2 in 2022^11^ were subsequently carried out using library generation methods optimized to produce saturating point mutations at each position, each characterizing a smaller number (10^4^) of additional sequences. However, these studies are inherently unable to provide comprehensive information on all possible mutations (*e*.*g*., 20^200^ *≈* 10^260^ for a 200 residue protein). Furthermore, the reported fitness estimate for any single sequence is noisy, impeding the isolation of important mutants and hampering the understanding of individual mutations.

The ability to computationally predict the effect of any mutation on viral fitness would allow scientists to quickly isolate which mutations might be problematic in the future. However, this dependence is difficult to model. Due to the complex nature of protein sequence-function relationships, the effect of combined mutations cannot be accurately estimated by adding or averaging the effects of single mutations. ^12–15^ Machine learning (ML) has shown substantial success modeling complex interdependencies; ^16^ for example, ML has been transformative for molecular biology, with approaches such as Alphafold predicting sequence-structure relationships. ^17–21^ Similarly, biological foundation models based on protein language models (PLMs), such as ESM^18^ and ProT5, ^22^ have learned generalizable sequence–function relationships directly from massive protein sequence datasets. Additionally, co-evolutionary models, including EVE^23^ and GEMME,^24^ have leveraged information from multiple sequence alignments to infer mutational effects. More recently, hybrid approaches that integrate PLM, co-evolutionary or explicit structural representations have shown improvements for specialized protein prediction tasks. ^25–29^ While these approaches are designed for broad transferability, their generalization to highly specialized or out-of-distribution regimes remains variable,^30,31^ often necessitating domain-specific adaptation via fine-tuning.^32–35^ Furthermore, the substantial computational resources required for these large models may pose practical constraints in some applications.

For well-defined, task-specific applications, dedicated models trained on focused data can be advantageous. The development of specialized protein fitness predictors has greatly benefited from high-throughput data generated by multiplex assays of variant effect (MAVE),^36^ such as deep mutational scanning (DMS).^37^ For example, Gelman et al. demonstrated that neural networks can learn sequence-function relationships from DMS data, capture nonlinear effects, and guide protein design.^38^ Tareen et al. introduced MAVE-NN for modeling genotype-phenotype relationships from MAVE data using a combination of statistical and neural network methods.^39^ Faure et al. developed MoCHI, a flexible framework that combines neural networks with interpretable statistical metrics, and uniquely enables higherorder epistatic modeling and integration of multimodal phenotypic data.^40^

These approaches have been applied to learn SARS-CoV-2 RBD fitness. Taft et al. described an approach for “deep mutational learning,” which integrates DMS data to rationally design mutagenesis libraries targeting specific regions of interest.^15^ Machine learning classifiers trained on these combinatorial libraries enabled accurate characterization of RBD variants as binders or non-binders to ACE2 and various therapeutic antibodies. Importantly, they found that point mutations were not additive at higher edit distances from wild type (WT), and combinatorial libraries were critical to model performance. As noted by the authors, this study focuses only on combinatorial mutations of select sites known to mutate in variants of concern (VOC), limiting the predictive scope.

Many others have developed predictive models of SARS-CoV-2 spike fitness;^41–51^ however, very few have focused on the capacity for early prediction when minimal sequence and experimental data are available from early strains. ^52^ In this work, we train RBD fitness predictors with DMS libraries and evaluate their ability to predict the fitness of unseen sequences. Accuracy is characterized using held-out data from the DMS experiment used for training, data from fully held-out DMS experiments, and by sampling the fitness landscapes defined using the output of the fitness predictor as an energy function with Markov chain Monte Carlo (MCMC).

Models trained on a large combinatorial DMS library (10^5^) initiated from the WT sequence are found to accurately characterize trends in held-out data from the experiment used for training and accurately predict patterns present in held-out DMS experiments, including sequences with much greater edit distances than found in training, such as omicron strains. Models trained on smaller libraries focused on attaining saturating point mutagenesis showed much worse predictive capacity. Fitness landscape simulations are assessed by their the capacity to predict the evolution of mutations observed in the population. The resulting sequence profiles prioritize hotspot residue positions and amino acid substitutions in VOCs comparable to profiles identified by applying a fitness threshold to the combinatorial DMS library. Though diverse combinatorial sequences are generated by the simulations, open questions remain about how to translate these landscapes into epidemeological predictions. Collectively, these findings demonstrate the importance of training predictors on multi-mutation data and the capacity of ML to characterize higher-order mutations more accurately than inferring simple models from point mutations, but highlight difficulties with evaluating the performance of predictors across sequence space. We expect that these conclusions will be applicable to other mutation prediction tasks for evolving pathogens and may help guide the analysis of future DMS datasets.

## Results and Discussion

### Machine learning architectures predict RBD binding affinity

Prediction performance was first quantified by training and evaluating models on data extracted from the same DMS experiment. Multiple predictors were trained via least-squares regression to link RBD sequence to ACE2 binding affinity. The DMS dataset used for training^6^ was experimentally initialized at the wild type (WT) RBD sequence and contains 105,525 variants with corresponding relative *K*_D_ values reported as Δ log *K*_D_ relative to the WT such that values above zero bind better then WT.

Predictors included linear models, a fully connected neural network (multilayer perceptron, MLP), ^16^ a message passing graph neural network (MPGNN),^53,54^ a transformer ^55,56^ and gradient-boosted decision trees^57,58^ with and without dropout (DART); ^59^ a highly-performing subset of results are shown in (**Figure 1**). Among all searched models and hyperparameters (**Tables S1-S2**), a shallow MLP using a one-hot sequence embedding achieved the lowest mean absolute error (MAE) of 0.27 Δ log *K*_D_, outperforming linear, DART, transformer and a structure-based MPGNN with test MAEs of 0.93, 0.46, 0.29, and 0.38 Δ log *K*_D_, respectively. These errors were calculated using a randomly selected test set comprising 10% of the total data. A separate validation set of the same size was used for hyperparameter optimization, ensuring that the test set remained completely uninvolved during training.

**Figure 1.**
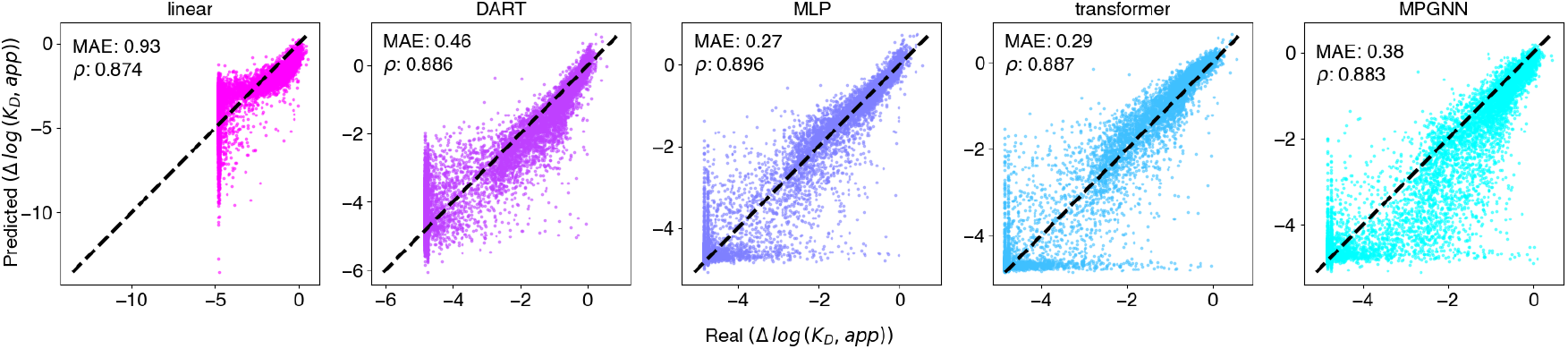
Summary of architectures’ performance on held out test set. Predicted versus experimentally measured Δ log *K*_D_ values are plotted with MAE and Spearman’s rank coefficients labeled on each plot.

An MPGNN with sufficiently expressive hyperparameters should ostensibly be able to outperform an MLP if defined via a suitable graph. However, similar to previous studies, ^44^ we do not see a gain in performance when using graph-based architectures. The tested MPGNNs were created using nearest neighbor relationships found in the WT structure, as representative structures are not known for most mutants, a possible avenue for improvement. However, since binding affinities are ensemble properties, static structural information may not be helpful as an inductive bias to predict mutational Δ log *K*_D_s in general.

The MLP predicts the fitness of higher-order mutations with far greater accuracy than simply adding or averaging the impact of point mutations. Though unsurprising, this highlights the importance of accounting for non-additive effects when analyzing DMS data, and the value of generating combinatorial mutation libraries.^12–15^ The MAE of the MLP remains low and relatively consistent regardless of the number of mutations, up to 9, the maximum number present in the training set (**Figure 2A**). However, the number of data points decreases with higher mutation counts, and despite maintaining low MAE, the Spearman’s rank correlation coefficient drops significantly with more than six WT mutations, likely because these mutants tend to all fall in the low binding, high noise region (**Figure 2B,C**). This illustrates, as previously noted,^60,61^ that validating the accuracy of variant effect predictors is a complex task. Differences in intended use can change which aspects of estimated performance are relevant; therefore, consideration of multiple metrics is critical.

**Figure 2.**
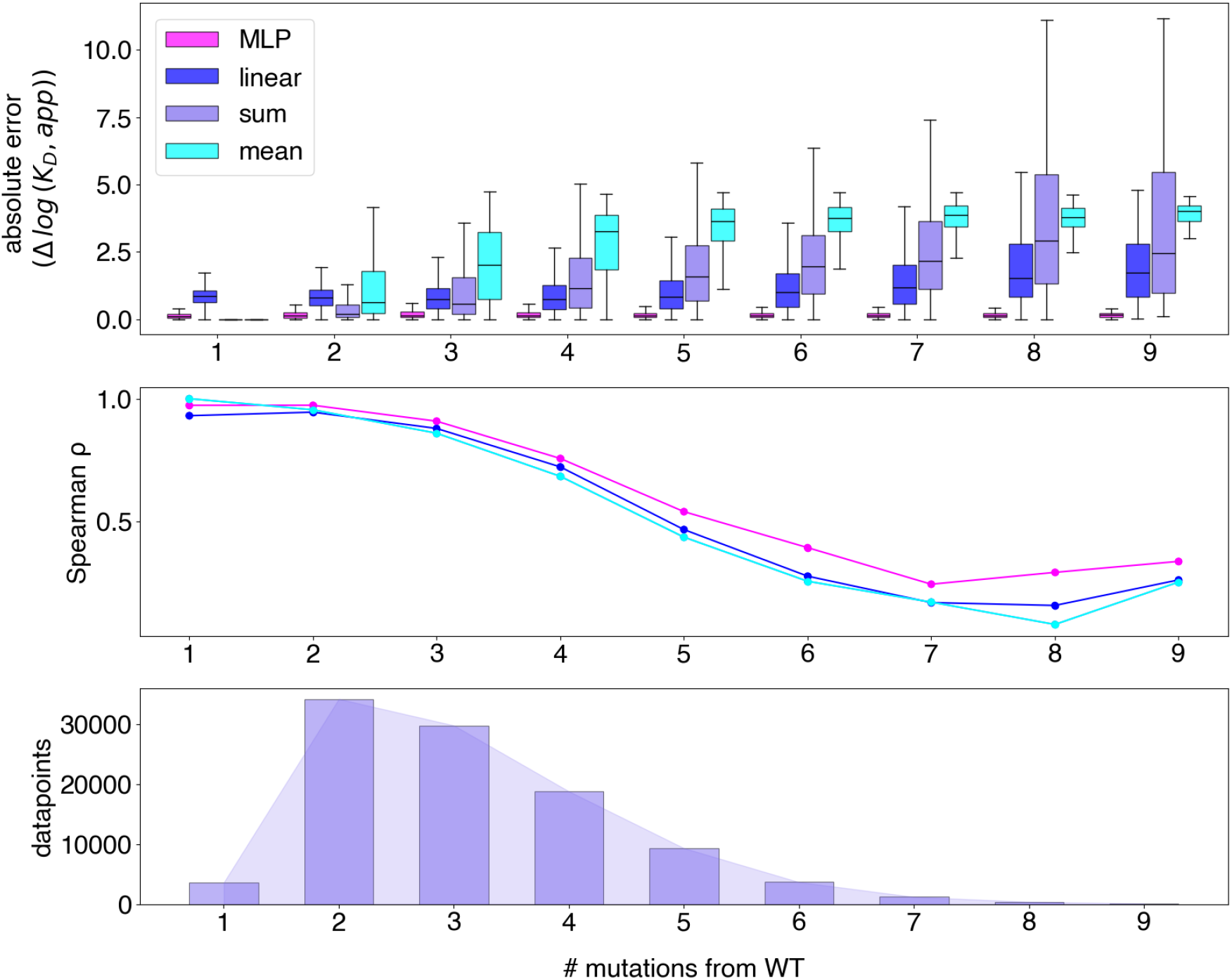
Predictions stratified by number of mutations from WT. A. box and whisker plots showing absolute error between true and predicted Δ log *K*_D_ for sequences in the library predicted with the MLP, linear model, or by taking the sum or mean of each point mutation in the sequence. B. Spearman’s rank correlation for each prediction in A. Note the sum and mean maintain the same rank and are therefore shown as a single cyan line. C. Amount of sequences in library containing each number of mutations from WT.

### Combining Multiple DMS Datasets

Several VOC out-competed the WT strain early in the pandemic. ^62^ As a result, subsequent RBD-ACE2 binding investigations were carried out with different starting sequences. In 2021 Bloom and colleagues published additional libraries initiated from N501Y, E484K, and B1.351 (N501Y, E484K, and K417N), ^10^ which we will refer to as the beta study, followed in 2022 by libraries initiated from omicron BA.1 and omicron BA.2 datasets,^11^ which we refer to as the omicron study. These binding measurements followed the design of the original WT study; however new library construction techniques aimed at efficiently saturating point mutations decreased the data available by an order of magnitude (*≈* 10^4^) per library. These new studies each included a control library constructed from the WT sequence, resulting in a total of seven new libraries in addition to the original study. Individual MLPs were trained on each library using the same method as above. Performance was evaluated on a random held out test set specific to each experiment. Models trained on these smaller datasets resulted in relatively low in-experiment performance with MAEs ranging 0.57 to 0.64 Δ log *K*_D_ compared to 0.27 Δ log *K*_D_ for models evaluated on the original dataset; Spearman coefficients ranged from 0.730 to 0.825 compared to 0.896 in the original. These results are from individual hyperparameter scans to obtain the optimal network architecture for each dataset (**Figure 3A**).

**Figure 3.**
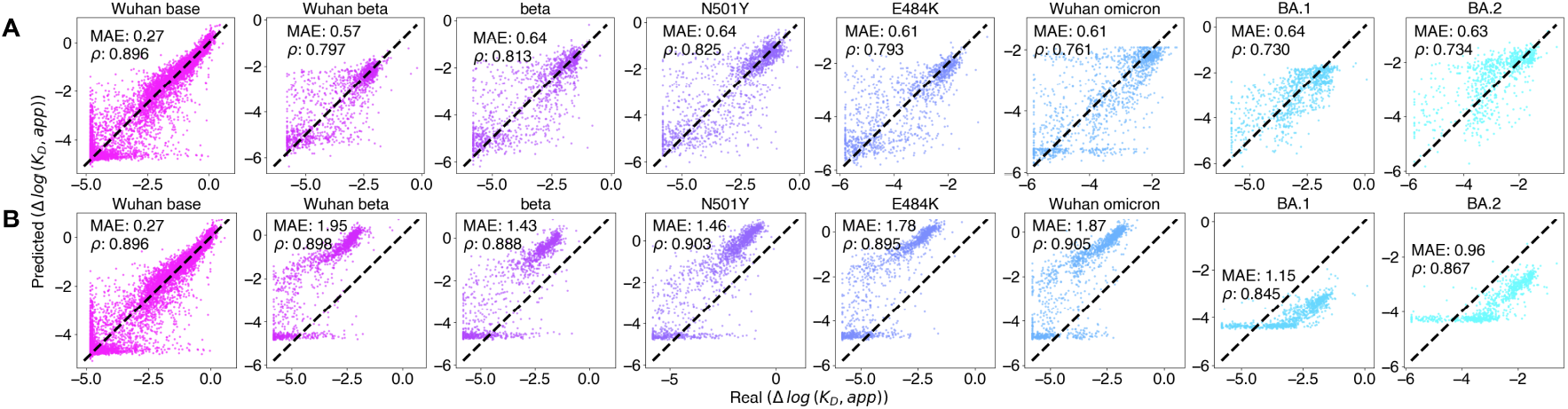
Summary of baseline extrapolation performance of initial model to 7 other DMS libraries. Each dataset is labeled based on the starting sequence used to generate library. These were conducted in 3 separate studies by Starr et al., each with a Wuhan (WT) reference sequence control. Studies include the original (base),^6^ B.1.351 (Wuhan beta, beta, N501Y, E484K), ^10^ and omicron (Wuhan omicron, BA.1, BA.2).^11^ A. Predicted versus real Δ log *K*_D_ values on for independently trained models. B. Predicted versus real Δ log *K*_D_ values of the original model on test sets from each individual library.

Since our model trained on the original WT library showed relatively low MAE up to nine mutations away and high Spearman coefficients up to five mutations away from the WT sequence (**Figure 2A,B**), we sought to determine how well this model trained only on the original WT library could predict data generated from independent experiments. The beta experiment contains libraries with up to seven mutations from WT and the omicron experiment contains libraries with between 14-19 mutations from WT. Surprisingly, the Spearman correlation coefficients of predictions with the original WT-trained model were higher than those found by training on data from the same experiment for every library, ranging 0.845 to 0.905. We attribute this primarily to the difference in training set size. However, these predictions corresponded to relatively high MAEs relative to the experiment-specific models with the original WT-trained model, ranging from 0.96 to 1.95 Δ log *K*_D_. Visual inspection suggests that these differing performance rankings are due to different experiment-specific offsets: the WT-trained model over-predicts affinity in all libraries besides omicron where affinity was under-predicted (**Figure 3B**). This prediction error may be due to difficulties in extrapolation or variations in experimental response curves. ^63^ These results suggest that predictors trained on one experiment may have the capacity to rank samples typical to heldout experiments, indicating a relatively high level of extrapolative ability. The improved performance characterizing samples within a given library can be useful for removing the noise typical to individual DMS observations; this application is explored further in the next section.

It is natural to ask whether the strong extrapolative performance of the WT-trained network on other smaller datasets can be improved by pooling training data from multiple experiments. First, fine-tuning the pre-trained WT network with the data drawn from the smaller experiments was attempted; this did not significantly improve performance on low-data experiments. Instead, training MLPs from scratch on combined data from all experiments proved more effective, resulting in a reduced MAE range of 0.38 - 0.46 Δ log *K*_D_ and increased Spearman correlation range to 0.893 - 0.907 for all datasets, except for those initiated from WT, which showed a *drop* in performance with 0.47 - 1.02 Δ log *K*_D_ MAE and 0.831 - 0.864 Spearman correlation (**Figure 4A**). The various libraries initiated from WT contain significant overlap in their sequences but contain different labels derived from each experimental setup. In order to account for the possibility of differing experiment response curves, a tuning head, which modifies the sequence-dependent prediction depending on the recorded experiment, was added to the MLP architecture.^63,64^ This decreased WT library MAEs to 0.28 - 0.41 Δ log *K*_D_, and increased the Spearman correlation range to 0.892 - 0.919. The non-WT libraries did not change substantially with this tuning (**Figure 4B**).

**Figure 4.**
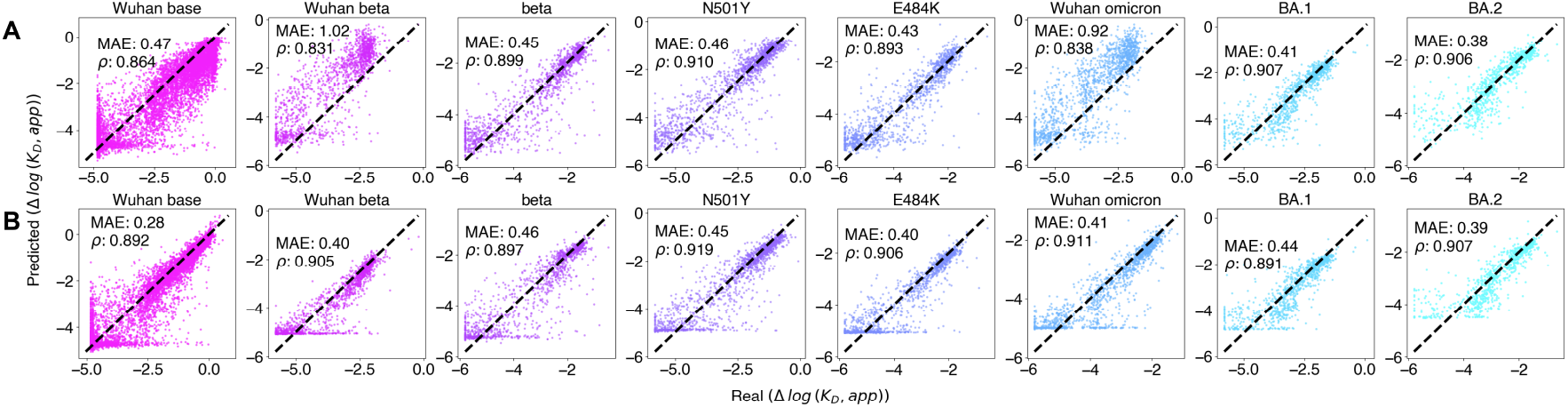
Summary of performance with aggregated training data from all DMS libraries. Each library is labeled based on the starting sequence as in Figure 3. Predicted versus real Δ log *K*_D_ values are plotted with MAE and Spearman’s rank coefficients shown for A. Without per-experiment tuning and B. With per-experiment tuning.

We note that per-experiment tuning may be useful for certain analyses, but it is not straightforward to utilize this model for the goal of predicting the outcome of held-out experiments as the corresponding response curve is not known. As demonstrated by the model without per-experiment adjustments, an MLP alone has the capacity to capture the outcome of multiple experiments when those experiments characterize distinct locations in sequence space. Ideally, the predictive accuracy of these various models would be validated by comparing to a “true” binding affinity that is not subject to variable experimental response curves; however, the authors are not away of a test set with this property. Instead, accuracy is evaluated based on the capacity to predict epidemiological trends in viral fitness, as described in the next section.

### Characterizing the fitness landscape

In order to link our trained predictors to observed population mutation trends, trained MLPs that accurately predicted RBD binding affinity from sequence were used to explore estimated fitness landscapes underpinning future VOCs using MCMC.^65^ Multiple definitions were used to create the fitness surface from the prediction of binding affinity. First, simulations aimed to assess the ability to predict VOC mutation trends solely from information on the WT sequence, and therefore the original WT-trained predictor was used. Since no knowledge of VOCs was present in the original DMS used for training, this represents a rigorous assessment of extrapolative performance.

First, we propose that an informative landscape may be defined by assuming the system is characterized by a Boltzmann-like distribution

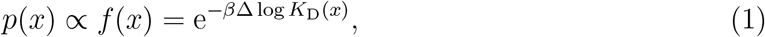

where *p*(*x*) is the probability of a particular sequence, *f* (*x*) is proportional to this probability, *β* is an unknown scalar, and Δ log *K*_D_ is the true change in binding affinity for a given mutant.

This definition suggests performing MCMC using the predicted output of a trained model in place of Δ log *K*_D_ with various values of *β*. As MCMC allows the system to explore sequences that are highly dissimilar to those present in the training data, simulations of this type represent a complex evaluation of the extrapolative abilities of a fitness predictor. Despite the promising out-of-experiment ranking performance, all simulations performed using Eq. (1) resulted in sequences with saturated mutagenesis (with every position containing a different amino acid compared to the WT) suggesting fundamentally limited model accuracy at high mutation levels (**Figure S1**). We also note that the combinatorial sequence space grows exponentially with sequence length; as a result, random mutation proposals are effectively biased toward highly mutated variants rather than few-site mutations.

Two modifications were made to produce more informative fitness simulations. First, to avoid excessive mutations driving sampling beyond mutation levels typical to the training data, sampled sequences were prohibited from exceeding 9 mutations relative to the the WT sequence. However, the resulting simulations still accumulated mutations up to the set maximum, with very little sampling below 7 mutations (**Figure S1**), suggesting that errors in fitness predictions may exist for specific mutants within 9 mutations of the WT. Therefore, to produce physically plausible mutation statistics and further focus sampling, we further modified the sampled distribution to be proportional to Eq. (2),

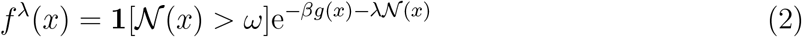

where *g* corresponds to the predicted affinity, *N* is the number of residues which match those present in the WT sequence, *λ* controls bias towards the WT sequence, **1** denotes the 0-1 indicator function, and *ω* places a limit on the maximum number of mutations allowed during the simulation. *λ* may be viewed as a tuning parameter controlling the average number of mutations found during the simulation. Eq. (2) is sampled by increasing the probability of proposing the WT amino acid at each position, while maintaining an equal probability of proposing non-WT amino acids, akin to the “favor native residue” function in Rosetta (**Figure S1**); this modified proposal mechanism controls the stationary distribution of the Markov chain (see Methods).

Eq. (2) corresponds to a minimally adjusted form of e^*−βg*(*x*)^ modified to obtain a given average mutation count. Similar to Eq. (1), corrections of this form may be rigorously derived using maximum entropy arguments^66,67^ and are used to incorporate experimental information into molecular dynamics force-fields.^68–70^ While corrections are often applied to modify simulations to match known experimental observations, here we use this approach to flexibly penalize mutations without having a precise reference value for the target average mutation and fitness level; instead, *β* and *λ* were set to result in mutational fluctuations spanning 0 to 9 to increase overlap with the training set, and an average model prediction which overlaps with 0 while extending to higher values (**Figure S1**).

Sequences sampled from *f*^*λ*^ were compared to the true sequence distribution observed in the human population deposited to Genbank as of Jan 31, 2025. These Genbank sequences represent total viral fitness, while the simulation distribution solely depends on predicted ACE2 binding affinity and sequence similarity to the wild type. Mutational frequencies present in the DMS data used for training were also compared against the Genbank data. As this DMS data contains random mutations that do not follow a fitness distribution dependent on ACE2 binding affinity, but rather reflect the experimental procedure used, samples were selected based on various Δ log *K*_D_ score cutoffs (**Figure S2**). An experimental Δ log *K*_D_ threshold of zero, corresponding to better or equal binding affinity compared to WT, was most predictive of VOC residue positions and mutations. Due to the large combinatorial sequence space, rather than focusing on exact sequence matches, we analyzed the propensity for mutation at each position. We also compared this to only analyzing greatest binding affinities from only the point mutation information, excluding combinatorial sequence information contained in the DMS dataset.

To determine if an ML predictor trained solely on DMS data initiated from WT could predict mutations in VOCs, we sampled the distribution given by the baseline model. We found 2.5 M steps sufficient to achieve consistent mutation profiles in three independent MCMC replicates (**Figure S4**). We compared the top 20 mutations predicted by the simulations to those found in Genbank as of January 31st 2025. Over 70% of the simulation-predicted mutations occur at VOC residues, including positions 452, 493, 498, and 501. In contrast, just over 30% of the score threshold-identified mutations from the WT DMS dataset overlap with VOC residues, including positions 339 and 452 (**Figure 5A**). The top scoring mutations out of the DMS point mutants only contained VOC positions 498, 501 and 505. Many residue positions are predicted to mutate to multiple amino acids. To account for this, we examined the top 20 residues with the highest mutation frequency, regardless of which amino acid they were mutated to, which revealed additional hotspot positions. The simulation additionally identified residue 373, while the DMS data highlighted positions 346, 493, and 501,and the point mutation only data identified 373, 477, and 484. In the top 5 positions, 4 out of 5 predicted by simulation overlap with VOC residues, compared to just 1 out of 5 from the DMS data and 2 out of 5 from the point mutations, although considering the top 20 positions results in the same total number of hotspots being predicted with each method(**Figure S3**). Collectively, this suggests that our simulations can enrich hotspot positions more effectively than analyzing the DMS data alone.

**Figure 5.**
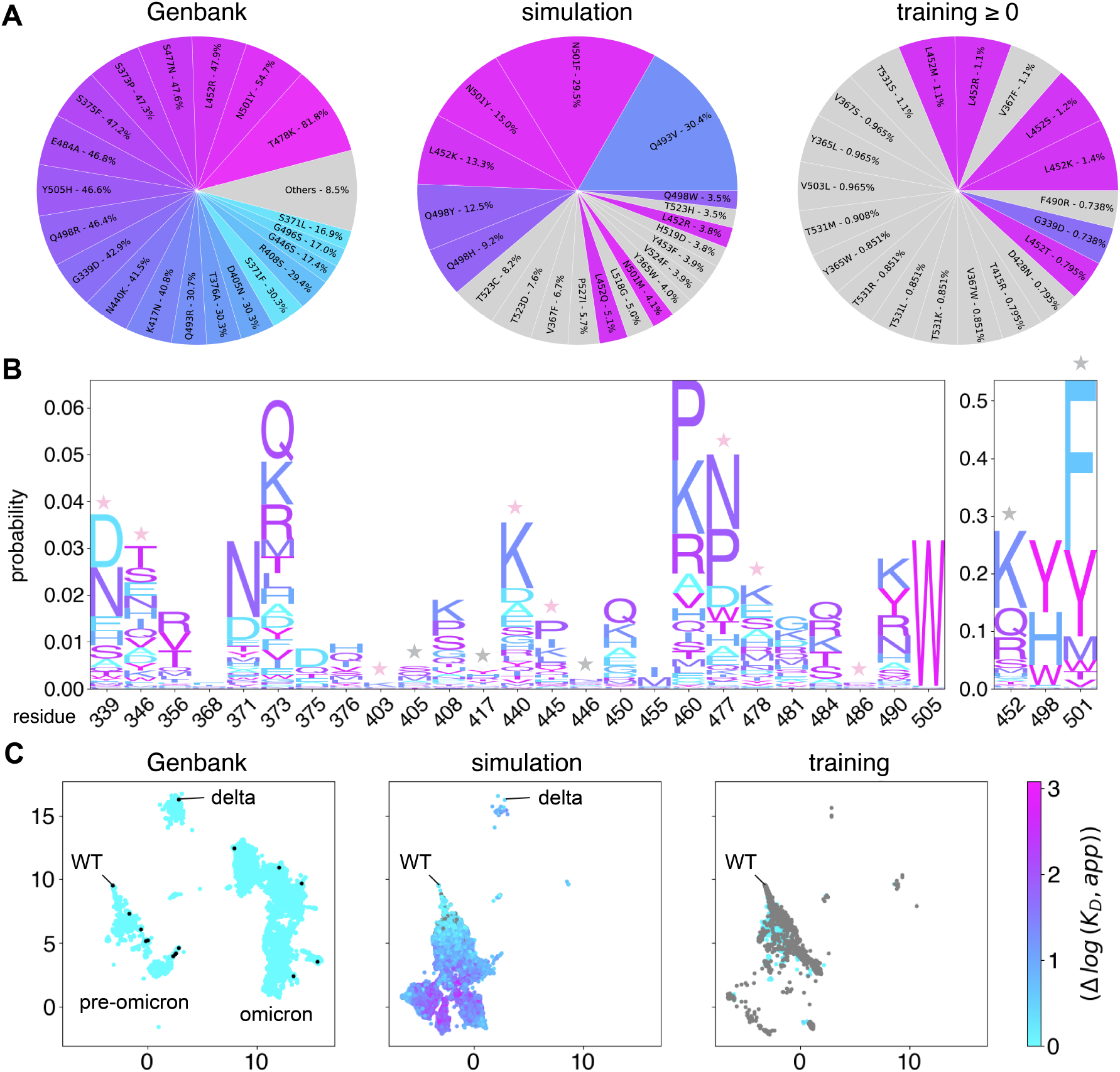
Sequence profile analysis. A. Pie charts showing top 20 mutations found in sequences from Genbank, simulation from WT with MLP trained on original WT data, and the training data sequences with scores greater than or equal to 0 Δ log *K*_D_ (better than WT). The percentage corresponds to the amount of total sequences in a given set have that mutation. Wedges are colored based on residue number, corresponding to those in the top 20 Genbank mutations. B. Logo plot of mutations in sequences from simulation for the 28 VOC residues. A pink star indicates the probable amino acid at a given position corresponds to a VOC mutation. A gray star indicates the most probable amino acid has the same physiochemical properties as the VOC mutation, such as polar, basic, and aromatic. C. UMAP of one-hot encoded sequences shown in A colored by Δ log *K*_D_. Unique Genbank sequences were colored with a score of zero, scores less than zero (ie. worse binders than WT) are colored gray. VOC from (Table 1) are shown as black dots on the first plot with representative variant categories labeled.

Next, the most frequent substitution at each position was analyzed for 28 mutations found in VOC strains (**Table 1**). Our simulation predicted the VOC amino acid as the most frequent mutation at nine positions: G339D, R346T, R403K, N440K, V445P, N460K, S477N, T478K, and F486P (**Figure 5B**). All of the correctly predicted mutations aside from S477N and T478K are specific to omicron strains. Additionally, at several other positions, the simulation did not predict the mutation found in VOC as the most frequent, however it predicted an amino acid with similar physiochemical properties. For example, N501 was often predicted to mutate to F instead of Y and L452 to K rather than R. Comparing this to the DMS or point mutation data, the same number of mutations was predicted, however their identities differed. Mutations R346T, S477N and F486P were predicted by the simulation but not thresholding, and L368I, N481K, and N501Y were identified with thresholding but not the simulation. Analysis of the highest scoring point mutants identified F486P, which was found in the simulation but not thresholding, but did not identify T478K which was found in both. Similar to the simulation, point mutation at position N501 to F has a higher affinity than to Y. This analysis suggests comparable performance for predicting the most frequent substitution for VOC residues with the simulation and the analysis of DMS data alone, though some mutations were only predicted by the simulation, while others were identified with the DMS but not the simulation.

**Table 1:** Summary of Variants of concern. Mutations predicted by the simulation are shown in bold. Omicron variants names are also labeled with Nextstrain clade scheme. ^71^.

| Name | Strain | RBD mutations |
| --- | --- | --- |
| Alpha | B.1.1.7 | E484K, **N501Y** |
| Beta | B.1.351 | K417N, E484K, **N501Y** |
| Gamma | P1 | K417T, E484K, **N501Y** |
| Delta | B.1.617.2 | **L452R**, **T478K** |
| Epsilon | B.1.42(7/9) | **L452R** |
| Eta | B.1.525 | E484K |
| Iota | B.1.526 | E484K |
| Iota' | B.1.526 | **S477N** |
| Kappa | B.1.617.1 | **L452R**, E484Q |
| Zeta | P2 | E484K |
| Mu | B.1.621(.1) | **R346K**, E484K, **N501Y** |
| 21K Omicron | BA.1 | **G339D**, S371L, S373P, S375F, K417N, **N440K**, G446S, **S477N**, **T478K**, E484A, **Q493R**, G496S, **Q498R**, **N501Y**, **Y505H** |
| 21L Omicron | BA.2 | **G339D**, S371F, S373P, S375F, T376A, D405N, R408S, K417N, **N440K**, **S477N**, **T478K**, E484A, **Q493R**, **Q498R**, **N501Y**, **Y505H** |
| 22A/B Omicron | BA.4/5 | **G339D**, S371F, S373P, S375F, T376A, D405N, R408S, K417N, **N440K**, **L452R**, **S477N**, **T478K**, E484A, **F486V**, **Q498R**, **N501Y**, **Y505H** |
| 23A Omicron | XBB.1.5 | **G339H**, **R346T**, L368I, S371F, S373P, S375F, T376A, D405N, R408S, K417N, **N440K**, **V445P**, G446S, **N460K**, **S477N**, **T478K**, E484A, **F486P**, F490S, **R493Q**, **Q498R**, **N501Y**, **Y505H** |
| 24A Omicron | JN.1 | **G339H**, K356T, S371F, S373P, S375F, T376A, **R403K**, D405N, R408S, K417N, **N440K**, **V445H**, G446S, N450D, **L452R**, L455S, **N460K**, **S477N**, **T478K**, N481K, V483-, E484K, **F486P**, **Q498R**, **N501Y**, **Y505H** |

As the models in this article were were trained to minimize error on held out data, their predictions may remove noise intrinsic to DMS measurements. This raises the question: can replacing experimental observations with predictions improve thresholding on DMS datasets? To assess whether this could improve prediction of VOC mutations, we performed inference on the training set and repeated analysis using the resulting Δ log *K*_D_ values for score thresholding. This resulted in a similar hotspot prediction to the original thresholding (**Figure S6**), indicating the simulation still performs better hotspot prediction. However, when considering the most frequent amino acid at the select VOC positions, the “denoised” score threshold resulted in a sequence profile with 13 of the most frequent substitutions correctly overlapped with the VOC mutations, including those identified by the simulation and experimental score threshold except for G339D and T478K. Additional mutations S371L, R408S, G446S, Q498R were identified with the denoised score threshold, but not by the simulation, experimental score threshold, or point mutation analysis (**Figure S5**). Though the simulation performed similarly to the DMS data with a threshold, the ML model-denoised thresholding of DMS data performed best at the task of retrospectively predicting the most probable mutation at a given position.

Simulations were next initiated from WT using the model trained on all DMS libraries using per-experiment tuning. Since these libraries contain information on beta and omicron BA.1 and BA.2 strains, predictions were assessed by focusing on 12 of the 28 mutations which appeared after BA.2 including BA.4/5, XBB.1.5 and JN.1 (**Table 1**). However, neither simulation of the multi-experiment model or thresholding of the corresponding DMS data identified new hotspots or substitutions not present in simulations performed by the original model. Both the original and aggregated training data identified the post-BA.2 residue 452 as most frequently mutated, while the original data also identified residue 490. Considering positions rather than frequent mutations, both additionally identify residue 346. Residues 452 and 460 are in the top 20 of both simulations trained on either model, with 460 representing another hotspot predicted by the simulations but not the DMS alone (**Figure S7**). The most frequent substitution at each position was the same for the thresholded training data and nearly the same for the simulation, however the simulation from the model trained on the aggregate data did not predict R346T. The denoising was not as helpful with this model either, predicting the same substitution but excluding L368I, V445P, N481K, F486P, and predicting L452 mutation to K instead of R (**Figure S8**). This suggests that inclusion of additional DMS libraries did not improve prediction of the most frequent substitution and worsened hotspot prediction. This may be attributed to difference in library construction techniques leading to smaller distributions around the starting sequence or differences in experimental response offsets; however, the exact cause is not clear. Centering the simulation bias around the BA.1 sequence did not result in additional hotspot residue or VOC mutation predictions; simulations with either model were less predictive than the simulations with the model trained on the original dataset and initiated from the WT sequence (**Figure S9**).

The combination of strong prediction accuracy on DMS datasets combined with the tendency of unbiased simulations to quickly accumulate saturating mutagenesis suggests while these predictors contain substantial information about parts of sequence space, certain regions uncharacterized in the training data create substantial challenges for simulation. While the introduction of additional DMS training datasets increases this coverage, they did not result in an increased prediction of VOC. This increase in data is dwarfed by the combinatorial growth in sequence space; for example, the omicron dataset provides 10^4^ additional data points; however, there exist more than 10^18^ possible mutants within its characteristic edit distance to the WT. Furthermore, as demonstrated by DMS denoising using a predictor trained with multiple datasets, improved performance on MAE across datasets may not translate to better downstream predictions of relevant mutations. Despite these challenges, simulations which generate informative mutations can be produced by biasing sampling to higher confidence areas of sequence space.

In order to better understand the overlap between the simulations, training datasets, and experimentally observed sequences we used principal component analysis (PCA) followed by Uniform Manifold Approximation and Projection (UMAP) ^72^ to project one-hot encoded sequences in two dimensions. When dimensional reduction was trained using only the simulated and VOC sequences, clusters showed overlap with all VOCs (**Figure S10**) using the model trained only on the WT data. However, when dimensional reduction was repeated using a combined set of sequences from Genbank, DMS training set, and the simulation, VOCs clearly clustered into three distinct regions including pre-omicron, delta, and omicron variants in the Genbank sequences. The simulated sequences primarily appear to explore pre-omicron and delta sequences, in addition to new unexplored regions. The DMS score thresholded sequences only appear to represent the pre-omicron sequences (**Figure 5C**). The dependence of the apparent coverage of VOCs on the reference set used for dimensional reduction underscores the sensitivity of visualization methods such as UMAP to input data selection. Despite this variability, however, differences in coverage between the data used for training and that produced by simulation underscore the extent to which extrapolation drives simulation results. The lack of overlap between simulation and omicron sequences is not surprising due to the biases required to drive simulation accuracy. Accurately characterizing sequence landscapes remains an open area of research with a range of strategies available,^73–75^ representing a promising avenue for future investigation.

## Conclusion

We have demonstrated that a combination of supervised ML with DMS and MCMC is a viable approach for simulating fitness landscapes and extracting meaningful information on possible mutations. Using a multilayer perceptron, we achieved highly accurate predictions of the SARS-CoV-2 RBD binding to ACE2. Notably, these predictions outperformed additive or averaged estimates in capturing the effects of combinatorial mutations suggesting an understanding of epsistatic relationships. Through retrospective comparison of predictions to population data, this predictor was shown to approximate an informative fitness landscape when combined with a maximum-entropy bias, generating a large number of unique combinatorial RBD sequences, identifying hotspot residues, and predicting mutations present in VOC (including omicron clades) from models only trained on variants near the WT sequence. Though unique VOC mutations were identified, aggregate sequence profiles did not identify a greater number of VOC mutations compared to analysis of high fitness sequences in the DMS data alone, leaving an open question of how to best translate generation of sequences with high predicted fitness into concrete epidemiological predictions. One limitation of the current implementation of evolutionary simulations is the reduction of viral fitness to the single metric of ACE2 binding, an avenue for future improvement. We introduce the open source package Mavenets: https://github.com/SztainLab/mavenets for reproducing and extending these findings, laying the groundwork for development of improved evolutionary predictors which may include multiple fitness metrics, alternative sampling algorithms, and prospective variant testing. Though only applied here to the case of SARS-CoV-2, these methods have potential application to a broad range of MAVE experiments and the *in silico* evolution of viral, bacterial, cancer-related, and designed proteins.

## Methods

### Data preparation

The WT ACE2 – RBD binding DMS datasets was obtained from https://media.githubusercontent.com/media/jbloomlab/SARS-CoV-2-RBD_DMS/master/results/binding_Kds/binding_Kds.csv. Data without Δ log *K*_D_ values were removed. Sequences associated with multiple entries within and across libraries were replaced with their average value, resulting in 105,525 unique sequences with Δ log *K*_D_ values. Data was randomly split into 80%/10%/10% points for train, valid, and test, respectively; except for training of DART models where 5-fold cross validation was performed over the combination of these train and validation sets. Additional datasets were preprocessed in the same manner as primary data. Δ log *K*_D_ was calculated for additional datasets by subtracting the WT log *K*_D_ as measured in the original library.

All universal resource locators are provided in Table S3.

### Prediction models

#### Multilayer perceptron

A multilayer perceptron (MLP) implemented with PyTorch^76^ was trained on various sequence embeddings to predict Δ log *K*_D_. Networks were created with ReLU^77^ activation on all layers except the output layer, which used no activation function. The mean squared error (MSE) between predicted and ground truth Δ log *K*_D_ values was used as the loss function for training. A batch size of 32 with the AdamW optimizer ^78,79^ was used to train until the convergence was observed via an uptick in the validation loss. No learning schedule was used with the optimizer; however, the schedule free optimizer^80^ was attempted during early hyperparameter tuning but did not produce better results. Networks were characterized by considering hidden layers of the following sizes: 8, 16, 32, 64, 128, and 256. Networks with hidden depths of 0-3 were created by independently varying the size of each layer (a depth of 0 corresponds to the linear model). In addition, scanning was performed over learning rates 3 *×* 10^*−*4^ and 1 *×* 10^*−*4^ and weight decay was scanned over 5 *×* 10^*−*3^, 1 *×* 10^*−*3^, 5 *×* 10^*−*4^, and 1 *×* 10^*−*4^. These various architectural and hyperparameters were explored via a grid search. We note that many MLP hyperparameter choices performed close to the optimal selected architectures; as a result, we do not attempt to interpret the optimal hyperparameters found for any experiment. Once optimal architectural and hyperparameter details were determined, these settings were used to train 3 networks with differing random initializations; the best performing model instance on the validation set was then evaluated on the test set.

#### Sequence embedding

Multiple different sequence embeddings (i.e., featurizations) were used as input to the MLP during early architecture explorations. A one-hot embedding was utilized based on the 20 canonical amino acids. Alternatively, a glycan one-hot embedding was defined by adding an additional token to the one-hot embedding denoting glycosylated asparagine residues based on the Nx[S/T] sequence motif.^81^ A UniRep^82^ embedding was used with the TAPEtokenizer^83^ utilizing a 1900-dimensional hidden state layer. SeqVec embeddings generated via Bio Embeddings were additionally tested, ^84^ as were ProtTrans-Bert-BFD, ProtT5-XL-UniRef50, ^56^ and ESM1b^85^ embeddings with default settings. Significant performance loss was observed with alternative embeddings compared to one-hot sequence embeddings; as a result, they were not explored further.

Other modeling approaches used only the one-hot sequence embedding for testing, with the exception of DART models which utilized equivalent categorical variables based on residue type and the transformer which used a custom scheme for embeddings.

#### Tuning heads

Certain neural networks were trained with per-experiment tuning (**Figure 4B**). This was implemented by first transforming the sequence into a scalar value using an internal network and subsequently refining this scalar value using function parametrized by the experiment identity; this is described in Eq. (3), where *g* denotes the full predictive model, *h* refers to an internal network, and *m* denotes the the experimental adjustment.

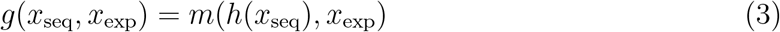

*x*_exp_ was defined to be an integer unique to each experiment and *x*_seq_ corresponds to the protein sequence. The internal network was set to be an MLP for the results shown in **Figure 4B**; results utilizing the transformer architecture yielded similar results (not shown). The experimental adjustment was defined to be a MLP with one hidden layer (input and output dimension of one) that is coupled to the output of *h* via a residual connection; ELU^86^ was used as the activation function to avoid dying neurons. The final (i.e., output) layer of this MLP is specific to a single experiment and other parameters are fully shared across experiments. In all cases hyperparameters of the per-experiment tuning head were optimized alongside those of the internal network. When training models using tuning heads, the first 20 training epochs used a linear combination of losses: the first loss corresponded to the mean squared error of the internal network and the second loss corresponded to the mean squared error of the experiment-adjusted prediction. Training was started weighting these two losses equally and linearly progressed to only penalizing the experiment-adjusted prediction by epoch 20; all remaining training was performed using only the experiment-tuned output. This scheduling was observed to stabilize training. The scanned sizes of the hidden layer were 1, 2, 4, 8, 16, 32, 64, 128, and 256.

### Message passing graph neural network

A message passing graph neural network was created using PyG^87,88^ to operate on a graph defined by a single structure of the WT RBD obtained from simulations performed by Barros et. al.^89^ Unless otherwise noted, training details mirrored those used for the MLPs. The structure was extracted from the trajectory by first clustering using cpptraj ^90^ via the hierarchical method with a random sieve of 10; this produced a total of 20 clusters. The average structure from the most populated cluster was then used to generate a residue graph with mdtraj.^91^ A one-hot residue representation of each sequence was used to describe the nodes, while edges were defined based on the distance between alpha carbons of each residue in the structure. In the optimal architecture, edges were connected based on a window in sequence space of 10 residues in the N and C terminal directions, as well as a distance cutoff of 15 Å in 3D space. Distances were featurized into 10 dimensional feature vectors using a Gaussian expansion and scaled with a factor of 75. Messages were passed 7 times, each containing an MLP with a 128 feature hidden layer. The following hyperparameters were scanned to identify the optimal architecture: window sizes: 5, 10, 15, and 20 Å; distance cut off: 10, 15, 20, 30 Å; distance feature vector dimensions: 0, 6, 8, 10, 12; scaling factor: 1, 10, 25, 50,75, 100.

### Transformer

A BERT-style^92^ (encoder-only) transformer was implemented in Pytorch using the SiLU activation function. ^93–95^ Unless otherwise noted, training details mirrored those used for the MLPs. The learning rate was set to 3 *×* 10^*−*4^. Embeddings were initialized by summing an embedding drawn from a look-up table based on residue type with an embedding vector specific to each position in the sequence. These embeddings are passed through multihead attention, layer norms, and an MLP as described in the pre-LN architecture.^96^ After MHA-based updates, each of the resulting embeddings is passed through a single shared MLP to produce a scalar value; this value is summed over the entire sequence to produce the model prediction. Hyperparameter optimization was performed scanning over the number of attention blocks, the number of attention heads, the embedding dimensionality, the dropout rate of the block MLPs, the dropout rate of the attention modules, the number of layers in the final MLP, and the dropout rate of the final MLP. A table of scanned values is found in **Table S2**.

### Multiple Additive Regression Trees with Dropout

Multiple Additive Regression Trees with Dropout (DART) ^59^ and gradient boosted decision tree models^57,58^ were created using the Light Gradient Boosted Machine library. ^97^ For brevity, both gradient boosted decision trees (without dropout) and Multiple Additive Regression Trees with Dropout are referred to as DART. The optimal model was found to use 511 leaves, dropout, 1399 trees, a drop rate of 0.3, a maximum drop of 50, minimum leaf data size of 1, a categorical l2 of 0, and a categorical smoothing of 5. Hyperparameters were scanned over using multiple local grid searches, scanning over the the presence of dropout (*i*.*e*., DART or gradient boosted decision trees), number of leaves, the rate of tree dropping, the presence of categorical smoothing and l2, the rate of tree dropping, and the minimum number of data allowed per leaf.

All hyperparameter combinations were trained for 5000 updates to determine the optimal number of trees. We note that a relatively large number of leaves was found to be beneficial to regression despite the low number of training points, likely implying a high degree of epistasis. Unlike other methods, hyperparameters were optimized using 5 fold cross validation performed on the the combined train and validation datasets used for other methods. All attempts treated inputs as categorical variables.

### Comparing combinatorial predictions

To compare higher-order mutation prediction with MLP versus adding or averaging the impact of point mutations, values of Δ log *K*_D_ for point mutations were obtained from the original WT dataset. This library contained 3602 unique point mutation values, corresponding to 89.6% coverage of saturating point mutations. Sequences in the library containing substitutions without a corresponding point mutation were removed from analysis. Sequences were stratified based on the number of mutations, and evaluated either with the MLP or by adding or averaging the Δ log *K*_D_ of each mutation present. We note that these models are distinct from the “linear” model in Fig. 1; this model was trained as a MLP with no hidden layer on a one-hot encoding.

### Monte Carlo Simulations

Markov Chain Monte Carlo (MCMC) simulations were performed to explore fitness landscapes implied by the output of trained models. For MCMC simulation a random position between 1 and 201 was selected for mutation to one of 20 canonical amino acids. The proposed mutant was evaluated by the model and accepted or rejected according to the Metropolis criteria.

MCMC performed using our trained model in place of Δ log *K*_D_ in Eq. (1) with Metropolis-Hasting^98^ moves leads to problematically high levels of mutations, likely driven by limited prediction accuracy on mutants far from those present in training set (SI). MCMC simulations are instead performed to target the following native-biased probability mass function

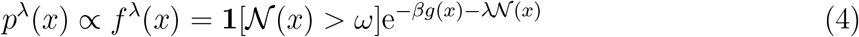

where *x* denotes a amino acid sequence, *N* denotes the number of residues in *x* which match those present in the WT sequence, *λ ∈* (*−*∞, ∞) determines a bias towards the WT sequence, **1** denotes the 0-1 indicator function, and *ω* places a limit on the maximum number of mutations allowed during the simulation. *ω* was set such that mutants with greater than 9 mutations were not considered. Several values for *β* were scanned and −10 was chosen to keep the center of the distribution of Δ log *K*_D_ outputs near 0, the fitness value of the WT. Values for *λ* were also tested and *−*2.5 was chosen to balance the diversity of the mutation landscape (Figure S4).

Eq. (4) is targeted by proposing single-residue mutations and accepting sequences using a modified form of the Metropolis procedure. Candidate 1-mutations were proposed using

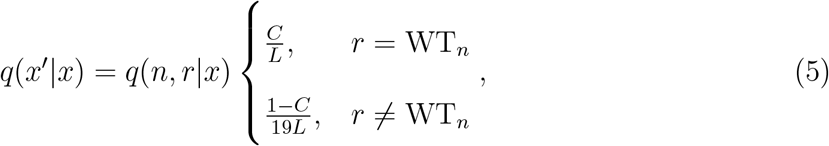

where *n* determines the index of the residue to mutate, *r* determines which amino acid the mutation will result in, and *L* denotes the number of mutable residues. Note that this a joint probability mass function over *n* and *r*; however, *n* does not appear as the corresponding marginal is uniform. Furthermore, note that this proposal distribution is asymmetric.

Proposed mutations are accepted according to a Metropolis criterion given by the following probability:

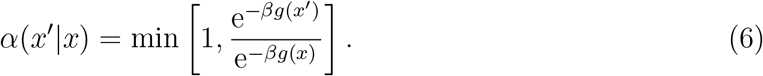

Application of Eqs. (5) and (6) result in the stationary distribution given by Eq. (4) when defining *λ* to be – ln 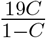 due to proposal asymmetry. The selected value for *λ* corresponds to *C* = 0.95.

In practice, sampling is performed by drawing a variate from a uniform distribution between 0 and 1 and comparing it to Eq. (6). If accepted, the mutant is set to be the current state of the MCMC simulation; if rejected, the state of the simulation is not changed. MCMC simulations were observed to return equivalent results on repeated runs, implying that simulations are well-converged.

### Genbank sequence collection

A total of 9,017,742 complete SARS-CoV-2 Receptor Binding Domain genome sequences were downloaded from NCBI Genbank ^99^ on Jan 31, 2025 by filtering for the SARS-CoV-2 taxon (taxid:2697049), human host, and a minimum length of 29,000 nucleotides. The 603bp RBD region (603 bp) from the Wuhan-Hu-1 reference genome (accession NC 045512.2,^100^ positions 22552–23155, NCBI RefSeq) served as the alignment target.

#### Sequence alignment and filtering

The pipeline used multiprocessing to parallelize alignment of the 257 GB dataset and sequences were loaded to a bounded queue to manage memory constraints. Alignments were performed with BioPython’s ^101^ PairwiseAligner with parameters mismatch score: −1 and gap opening penalty: −2. A windowed alignment strategy was employed during alignment focusing first on the expected RBD region with iterative expansion before falling back to full sequence alignment when necessary. Ambiguous nucleotides (N) replaced with their corresponding Wuhan reference nucleotides. The 603-nucleotide RBD coding sequences were converted to 201 amino acids using the standard genetic code table. Sequences exhibiting *>*20% divergence from reference in the RBD region were excluded (*<*500 sequences). The final dataset after alignment and filtering yielded 8,255,834 RBD sequences (91.55% of total), representing 27,980 unique RBD sequence variants consisting of combinations of 1,311 distinct mutations.

## Data availability

All code necessary for featurizing, training and evaluating models and for running MCMC simulations can be found at: https://github.com/SztainLab/mavenets. Code for reproducing all figures in the manuscript can be found at https://github.com/SztainLab/ML_evolution_SARS-CoV-2. Links to DMS data used can be found in (**Table S3**).

## Acknowledgement

We gratefully acknowledge funding from the European Commission (Grant No. ERC CoG 772230 “ScaleCell”),the Deutsche Forschungsgemeinschaft DFG (RTG2433 DAEDALAUs, SFB1114/C03, FOR2518, NO825/4). LR acknowledges funding through European Union’s Horizon 2020 research innovation programme under the Marie Sklodowska-Curie grant agreement No 897414, AD acknowledges funding through a Flagship Initiative Engineering Molecular Systems Postdoctoral Fellowship, and TS acknowledges startup funding from the University of Michigan Medicinal Chemistry Department. We thank Frank Noé for invaluable insights and discussions throughout this project. We also appreciate the technical assistance of Cornelius Hoffmann and Allen Bailey.

## Author contributions

TS conceived of the project. TS and AD wrote manuscript with contributions from all authors. Code for ML and analysis was developed collaboratively by all authors. AD consolidated and implemented these into the final software package.

## Supporting Information Available

Supplementary materials will be made available upon publication.

- Table S1. Optimal hyperparameters for MLPs used in this study.
- Table S2. Scanned Transformer values for the original WT dataset.
- Table S3. Summary of DMS datasets
- Figure S1. Optimiziation of MCMC simulation.
- Figure S2. VOC predictions with DMS training data using Δ log *K*_D_ score thresholds.
- Figure S3. Top 20 residue positions identified.
- Figure S4. Sequence profile of three MCMC replicates.
- Figure S5. Sequence profile of training data.
- Figure S6. Top 20 mutations with denoised score threshold.
- Figure S7. Top 20 mutations and positions colored by the 12 mutations appearing after BA.2.
- Figure S8. Sequence profiles of the 12 mutations appearing after BA.2.
- Figure S9. Top 20 mutations, positions, and sequence profiles from simulations centered around the BA.1 sequence.
- Figure S10. UMAP of one-hot encoded sequences trained only on simulation and VOC.

## TOC Graphic

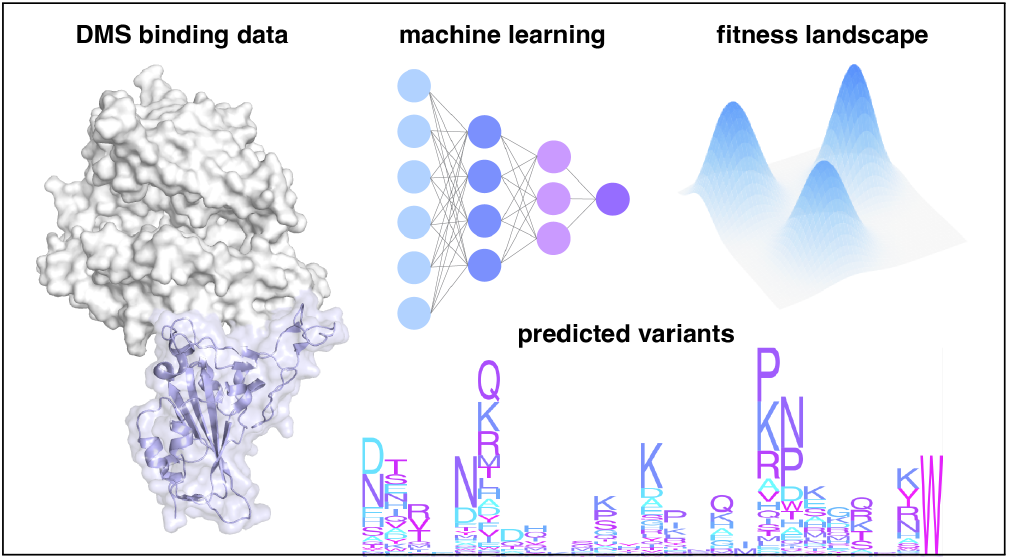

