## Supporting Information for "Machine Learning Driven Simulations of the SARS-CoV-2 Fitness Landscape from Deep Mutational Scanning Experiments"

Table S1: Optimal hyperparameters for MLPs used in this study

| model | layers | weight decay | learning rate |
|---|---|---|---|
| base WT | [16, 32, 16] | 0.0001 | 0.0001 |
| all data untuned | [8, 256, 32] | 0.0001 | 0.0001 |
| all data tuned | [8, 8, 8] | 0.0005 | 0.0001 |
| Wuhan - beta experiment | [16, 64] | 0.005 | 0.0003 |
| B.1.351 | [16, 64] | 0.0005 | 0.0003 |
| N501Y | [8, 8, 8] | 0.001 | 0.0003 |
| E484K | [8, 32, 64] | 0.001 | 0.0003 |
| Wuhan - omicron experiment | [8, 8, 16] | 0.005 | 0.0003 |
| BA.1 | [32, 8, 8] | 0.0005 | 0.0001 |
| BA.2 | [256, 256, 256] | 0.001 | 0.0001 |

Table S2: Scanned Transformer values for the original WT dataset. Optimal values shown in bold.

| Variable | Values |
|---|---|
| Num. Block | 1, 2, 3, 4, 5, **6** |
| Num. Heads | 2, 4, **8**, 16 |
| Embedding Size | 16, 32, **64** |
| MHA Dropout | **0.05**, 0.1, 0.2 |
| Block MLP Dropout | 0.05, **0.1**, 0.2 |
| Num. MLP Final Layers | 0, **1**, 2, 3 |
| Final MLP dropout | 0.0, 0.05, **0.1** |
| Weight Decay | **5e-3**, 1e-3, 5e-4 |

Table S3: Summary of DMS datasets.

| Dataset | Strains | Size | Source |
|---|---|---|---|
| Original | Wuhan Hu 1 | 105525 | link |
| B1.351 | Wuhan Hu 1 | 13,449 | link |
| | N501Y | 14682 | |
| | E484K | 11910 | |
| | B1351 (Beta: K417N, E484K, N501Y) | 12,722 | |
| Omicron | Wuhan Hu 1 | 13,892 | link |
| | BA1 | 9,161 | |
| | BA2 | 8,191 | |

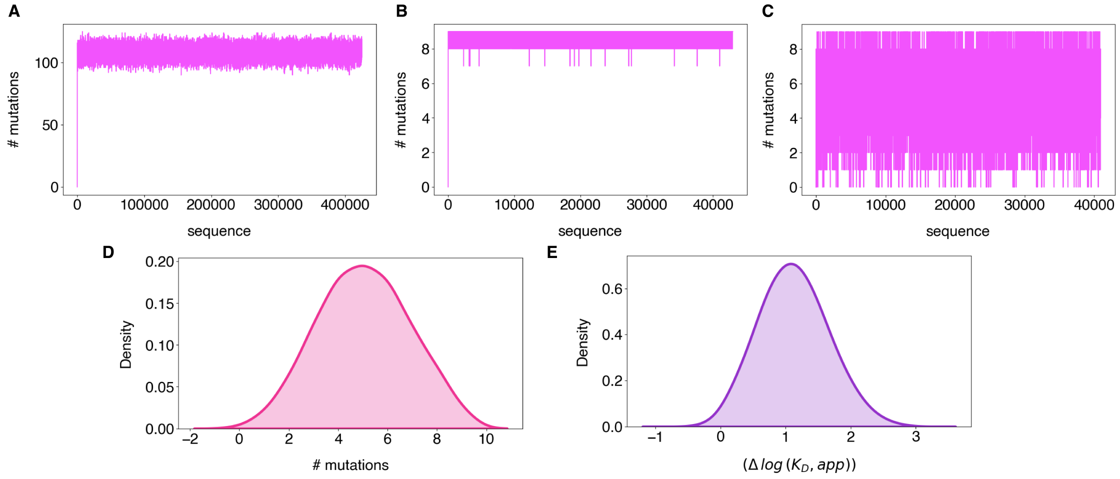


Figure S1: Optimization of MCMC simulation. A. Without any proposal bias. B. With maximum mutation count set. C. With proposal bias in Eq. 2 applied. D. Smoothed mutation distribution of simulation with optimal parameters E. Smoothed $\Delta \log K_{\mathrm{D}}$ distribution of simulation with optimal parameters.

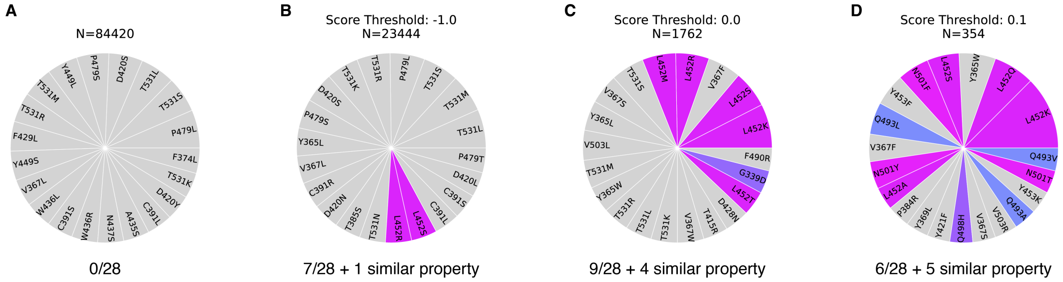


Figure S2: VOC predictions with DMS training data using $\Delta \log K_{\mathrm{D}}$ score thresholds. Score threshold and number of sequences which pass are labeled above each chart. Number of VOC correctly predicted as the most frequent mutation at a given position are indicated below each chart.

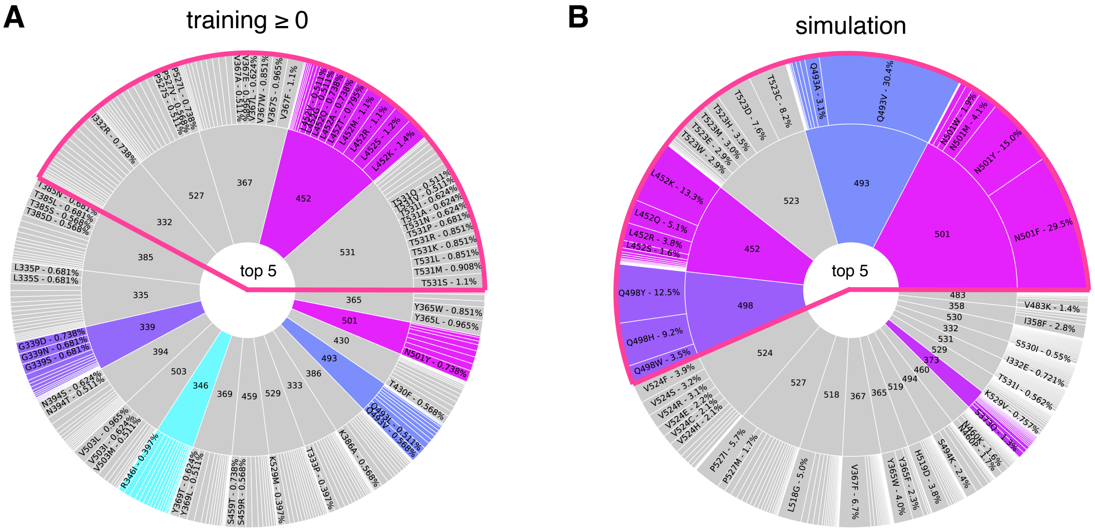


Figure S3: Top 20 residue positions identified. A. DMS training data using a $\Delta \log K_{\mathrm{D}}$ score threshold of 0 (better than WT) B. MCMC simulation. The top 5 residue positions are highlighted with a pink outline. Sequence profile is shown as a nested pie chart with the inner layer corresponding to the residue position, and outer layer indicating which amino acid substitutions are most frequent at a given position.

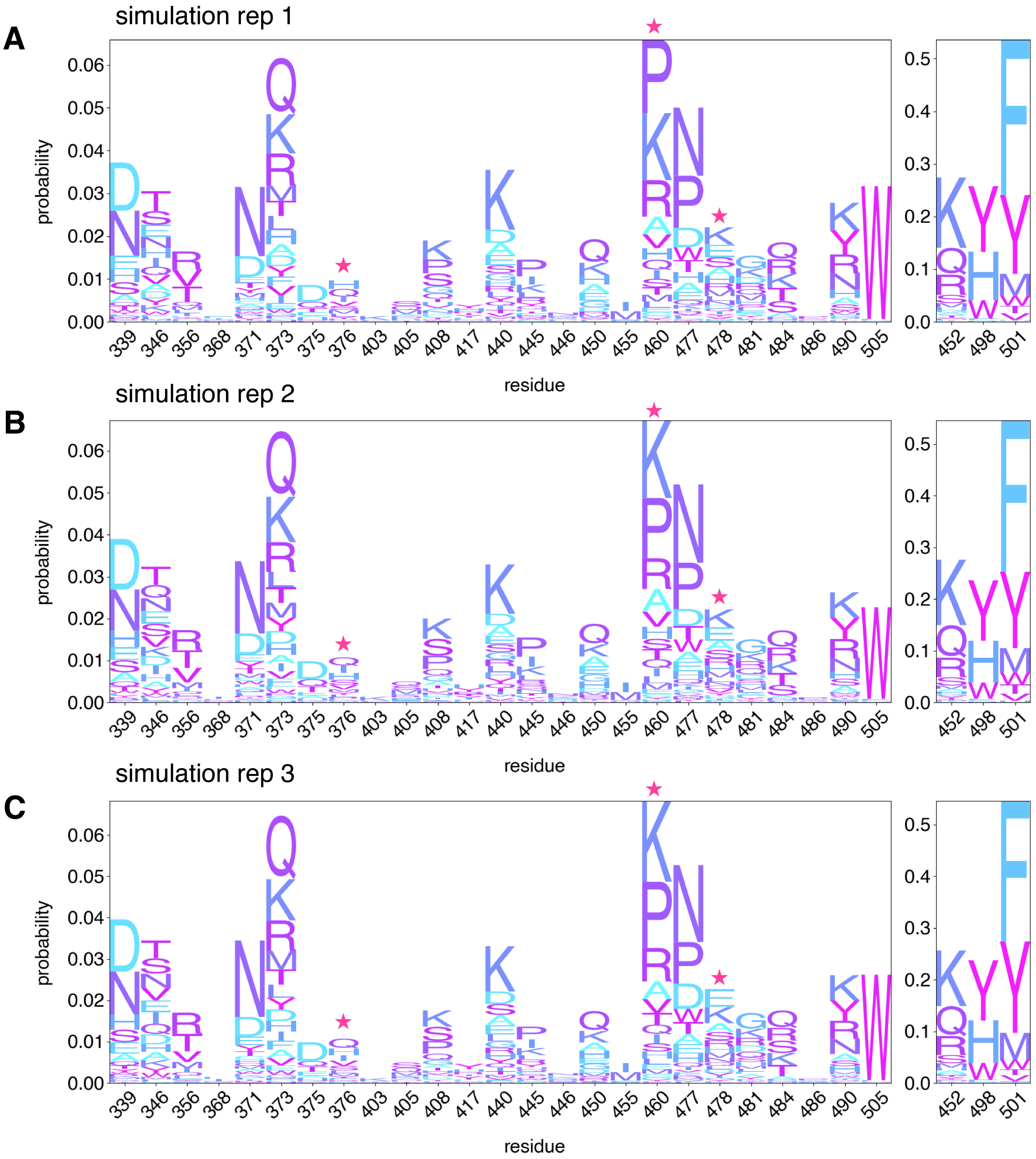


Figure S4: Sequence profile of three MCMC replicates. Residues with a difference in most prevalent amino acid mutation are highlighted with a pink star.

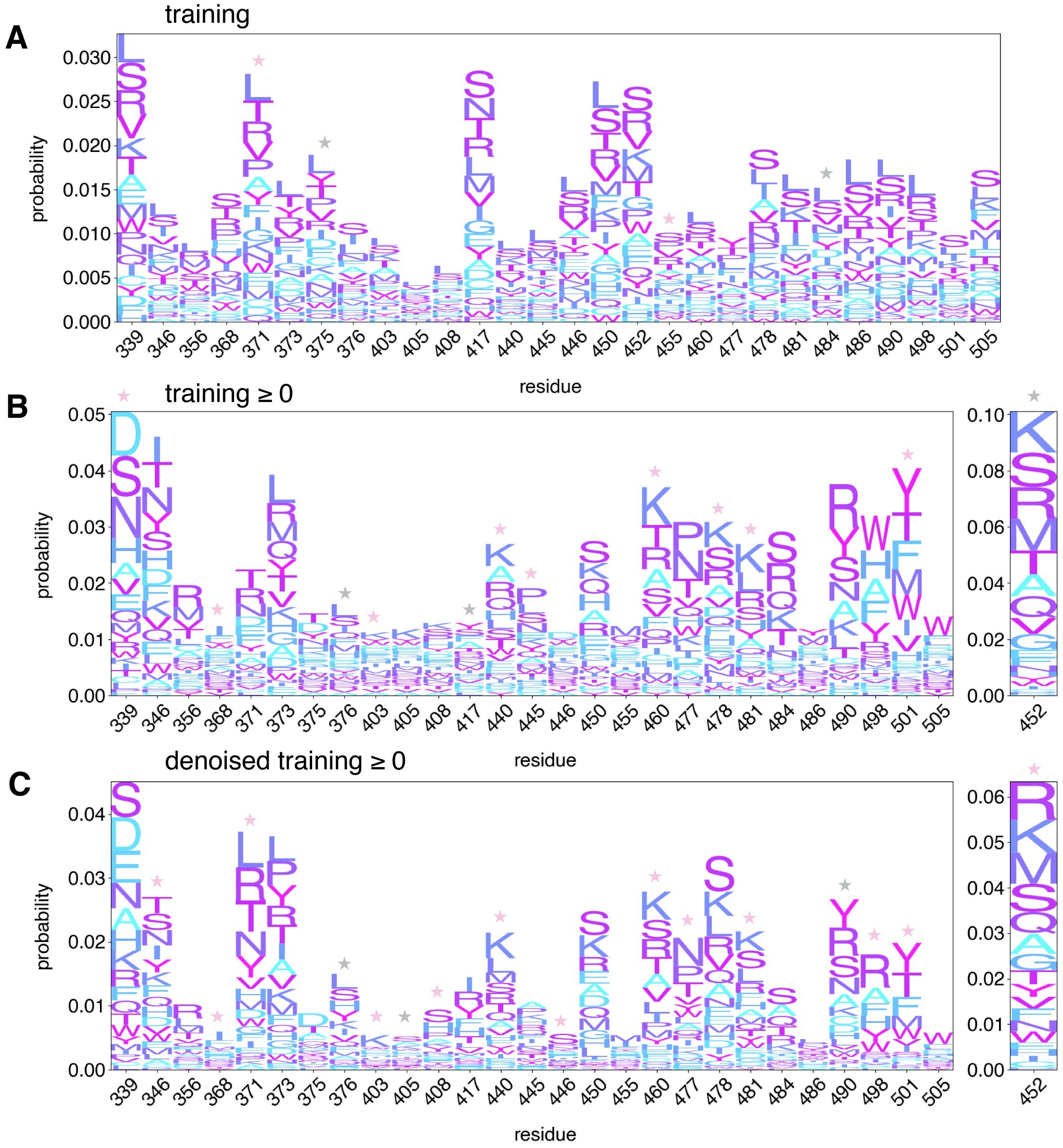


Figure S5: Sequence profile of training data. A. Without a score threshold B. With a score threshold of 0 (better than WT) and C. With a score threshold of 0, with scores recalculated from running sequences through the trained ML model. A pink star indicates the probable amino acid at a given position corresponds to a VOC mutation. A gray star indicates the most probable amino acid has the same physiochemical properties as the VOC mutation, such as polar, basic, and aromatic

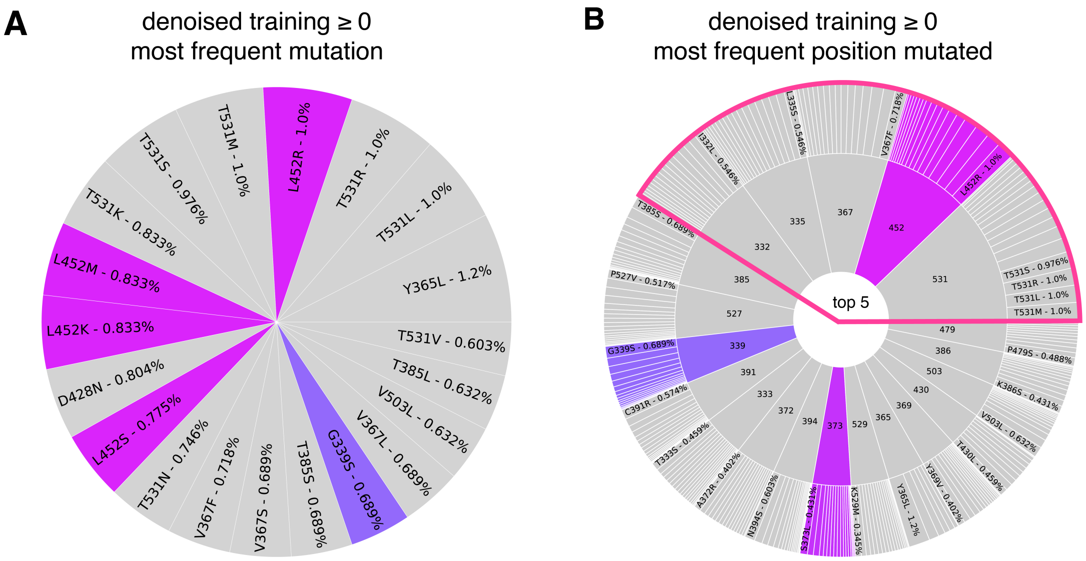


Figure S6: Top 20 mutations with denoised score threshold. A. Top 20 mutations overall, B. Top 20 positions that are mutated, regardless of which amino acid they are mutated to. Outer ring slices indicate different amino acids with the most frequent substitution labeled.

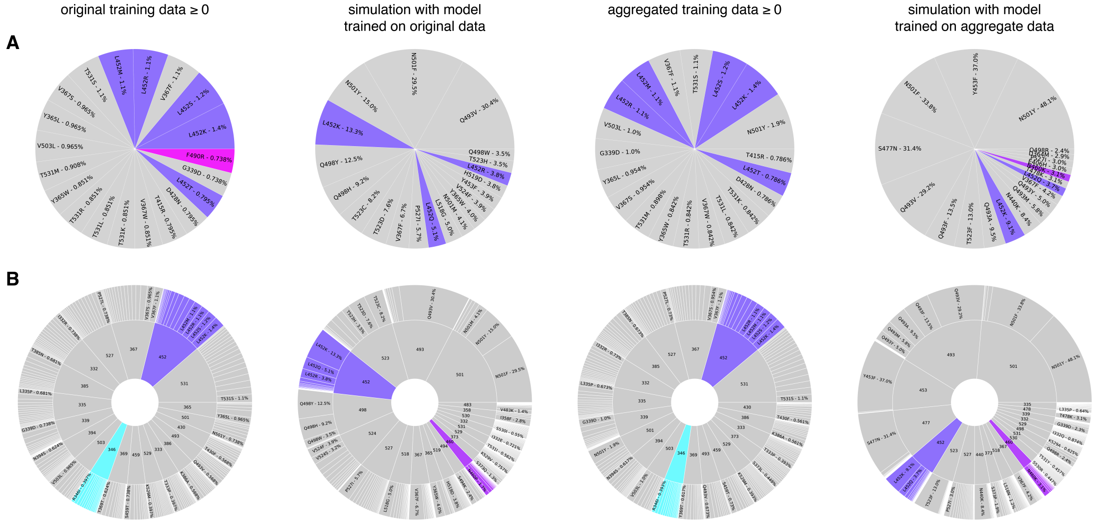


Figure S7: Top 20 mutations and positions colored by the 12 mutations appearing after BA.2. (See **Figure S8** for the 12 residues considered). A. Top 20 mutations overall from the original training data with threshold, simulation from model trained on original data, aggregated training data from all 8 DMS libraries with threshold, and simulation from per-experiment tuned model trained on data from all 8 libraries. B. Top 20 position regardless of mutation for each. The outer ring slices indicating different amino acids, with the most frequent substitution labeled.

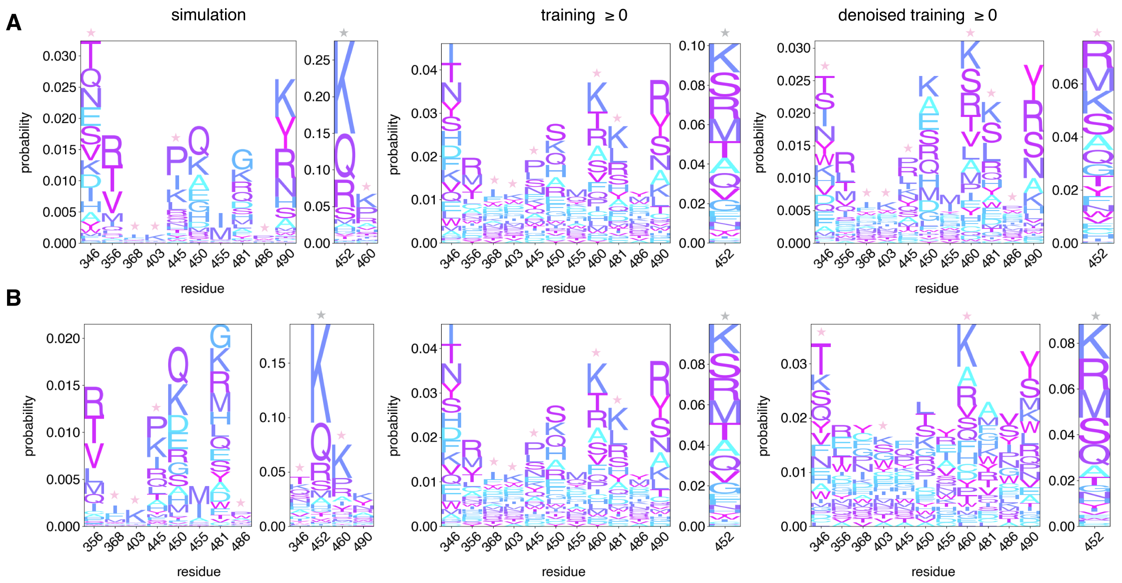


Figure S8: Sequence profiles of the 12 mutations appearing after BA.2. A. profiles from the simulation using model trained on the original data, the original training data with a $\Delta \log K_{\mathrm{D}}$ threshold of 0, and denoised training data, with the threshold recalculated from running on the training data with the model trained on the original data. B. Sequence profiles for each category in A. using the per-experiment tuned model and training data from aggregating all 8 DMS libraries. Note the sequence profiles of the training data with threshold of 0 for the original and aggregate data appear identical because only a small number of additional sequences from the new libraries passed the score threshold. Various threshold values were considered and a threshold of 0 was still the most predictive. A pink star indicates the probable amino acid at a given position corresponds to a VOC mutation. A gray star indicates the most probable amino acid has the same physiochemical properties as the VOC mutation, (ie. L452R predicted as K is also a mutation from non-polar to basic amino acid).

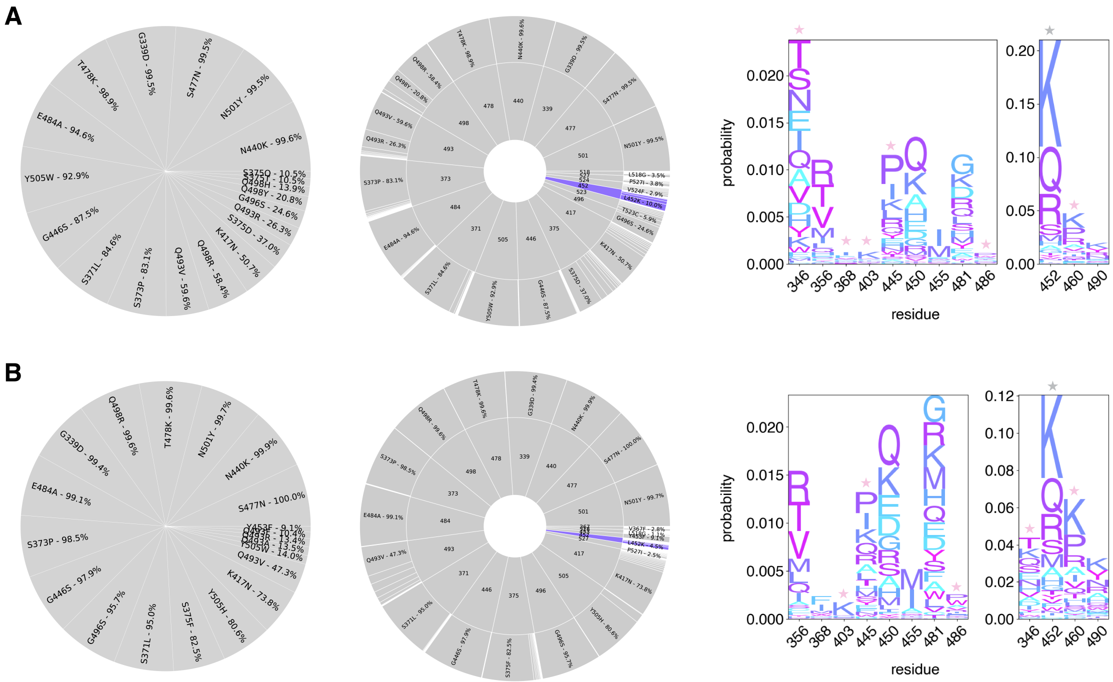


Figure S9: Top 20 mutations, positions, and sequence profiles from simulations centered around the BA.1 sequence. A. Simulations using model trained on original DMS data. B. Simulations using model trained on aggregated data with per-experiment tuning. Residues are colored in pie charts if they are in the list of 12 mutations appearing after BA.2. All charts follow the same formatting detailed in the previous figures.

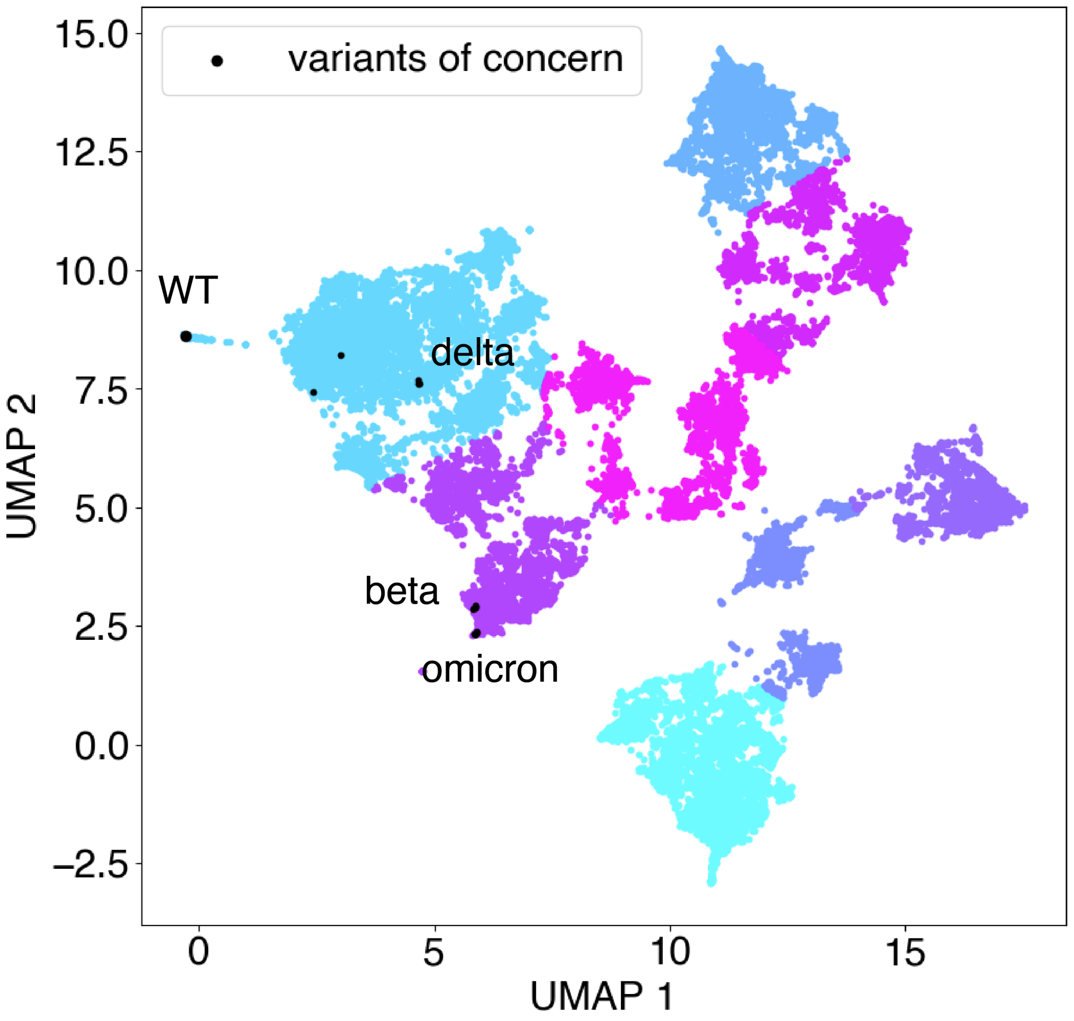


Figure S10: UMAP of one-hot encoded sequences trained only on simulation and VOC. Simulation results are from original WT-trained model. Colors are based on K-means clustering. VOCs from **Table 1** are plotted as black dots with select categories labeled.
